# Genetic variation in herbivore resistance within a strawberry crop wild relative (*Fragaria vesca* L.)

**DOI:** 10.1101/728360

**Authors:** Daniela Weber, Paul A. Egan, Anne Muola, Johan A. Stenberg

## Abstract

To decrease the dependency on chemical pesticides, the resistance of cultivated strawberry to pests needs to be increased. While genetic resources within domesticated varieties are limited, wild genotypes are predicted to show high heritable variation in useful resistance traits. We collected 86 wild accessions of *Fragaria vesca* L. from central Sweden and screened this germplasm for antibiosis (pest survival and performance) and antixenosis (pest preference) traits active against the strawberry leaf beetle (*Galerucella tenella* L.). First, extensive common garden experiments were used to study antibiosis traits in the sampled plant genotypes. Heritable genetic variation among plant genotypes was found for several antibiosis traits. Second, controlled cafeteria experiments were used to test for plant genetic variation in antixenosis traits. The leaf beetles avoided egg laying on plant genotypes possessing high antibiosis. This indicates a high degree of concordance between antibiosis and antixenosis, and that the beetles’ egg-laying behaviour optimizes the fitness of their offspring. The existence of high genetic variation in key resistance traits suggests that wild woodland strawberry contains untapped resources that are sought to reduce pesticide-dependence in cultivated strawberry. Given that only a very small portion of the species’ distribution area was sampled, even higher variation may be expected at the continental scale. As a whole, the genetic resources identified in this study serve to strengthen the position of woodland strawberry as a key crop wild relative.

## Introduction

Higher intrinsic resistance in crop plants against pests is a promising option to gain independence from heavy pesticide use during cultivation (Broekgaarden et al. 2011; Smith and Clement 2012). Especially in cultivated strawberry, the demand for alternative pest control approaches is growing, since the fruits are amongst the food with the highest pesticide residues (Fernandes et al. 2011; Godfray et al 2014; Parikka and Tuovinen 2014). Genetic resources for resistance against pests are limited within current strawberry cultivars and breeding programs (Liston et al. 2014; Chen et al 2015). A good source to enhance traits for pest resistance is the naturally existing variation among the crop wild relatives (CWR) of cultivated strawberries (Egan et al. 2018). Known as ‘rewilding’ or ‘inverse breeding’, beneficial traits such as plant resistance to herbivores can be restored in future varieties by including CWR germplasm in breeding programs (Broekgaarden et al. 2011; Andersen et al. 2015; Palmgren et al. 2015; Dempewolf et al. 2017). For example, a resistance trait to control the cabbage root fly *Delia radicum* L. (Diptera: Anthomyiidae) was identified in wild white mustard (*Sinapis alba* L.), and successfully integrated in oilseed rape (*Brassica napus* L.) and rutabaga (*B. napus. var. napobrassica*) (Ekuere et al. 2005; Malchev et al. 2010).

In recent times, woodland strawberry (*Fragaria vesca*) has been recognized as a model species and potential source of diversity for *Fragaria* breeding programs, including those for the garden strawberry (*Fragaria* × *ananassa*) (Longhi et al. 2014; Urrutia et al. 2015; Egan et al. 2018). Owing to the large geographic distribution of the wild *F. vesca* (Hilmarsson et al. 2017) and its natural adaptation to a range of habitats, the resulting genetic diversity of *F. vesca* is a promising source of desirable traits for abiotic and biotic stresses (Amil-Ruiz et al. 2011; Liston et al. 2014). The sequenced genome of *F. vesca* shares a very high degree of synteny with the hybrid *F. × ananassa* (Tennessen et al. 2014) and has already permitted greater insight into the molecular mechanisms underpinning flowering (Mouhu et al. 2013; Hollender et al 2014) and fruiting (Kang et al. 2013). In addition, *F. vesca* itself represents a valuable niche crop with a growing market due to the intense taste of its fruits (Ulrich et al. 2007; Doumett et al. 2011). However, knowledge about resistance against herbivores in *F. vesca* is still scarce (Muola et al. 2017), and, thus, further exploration of resistance in wild germplasm is needed to fully utilize the available gene pool.

We used wild *Fragaria vesca* originating from Sweden to investigate genetic variation in resistance to a chewing herbivore, the strawberry leaf beetle *Galerucella tenella* (L.) (Coleoptera: Chrysomelidae)(Fig. 1). *Galerucella tenella* is a pest insect in open strawberry plantations in Russia and Northern Europe, which is commonly controlled by spraying with insecticides (Parikka & Tuovinen 2014; Stenberg 2014). Larvae and adult beetles feed on strawberry leaves as well as on the flowers, which reduces pollination success, and larvae can furthermore damage fruits (Olofsson & Pettersson 1992; Parikka & Tuovinen 2014, Muola et al. 2017).

**Fig. 1.**
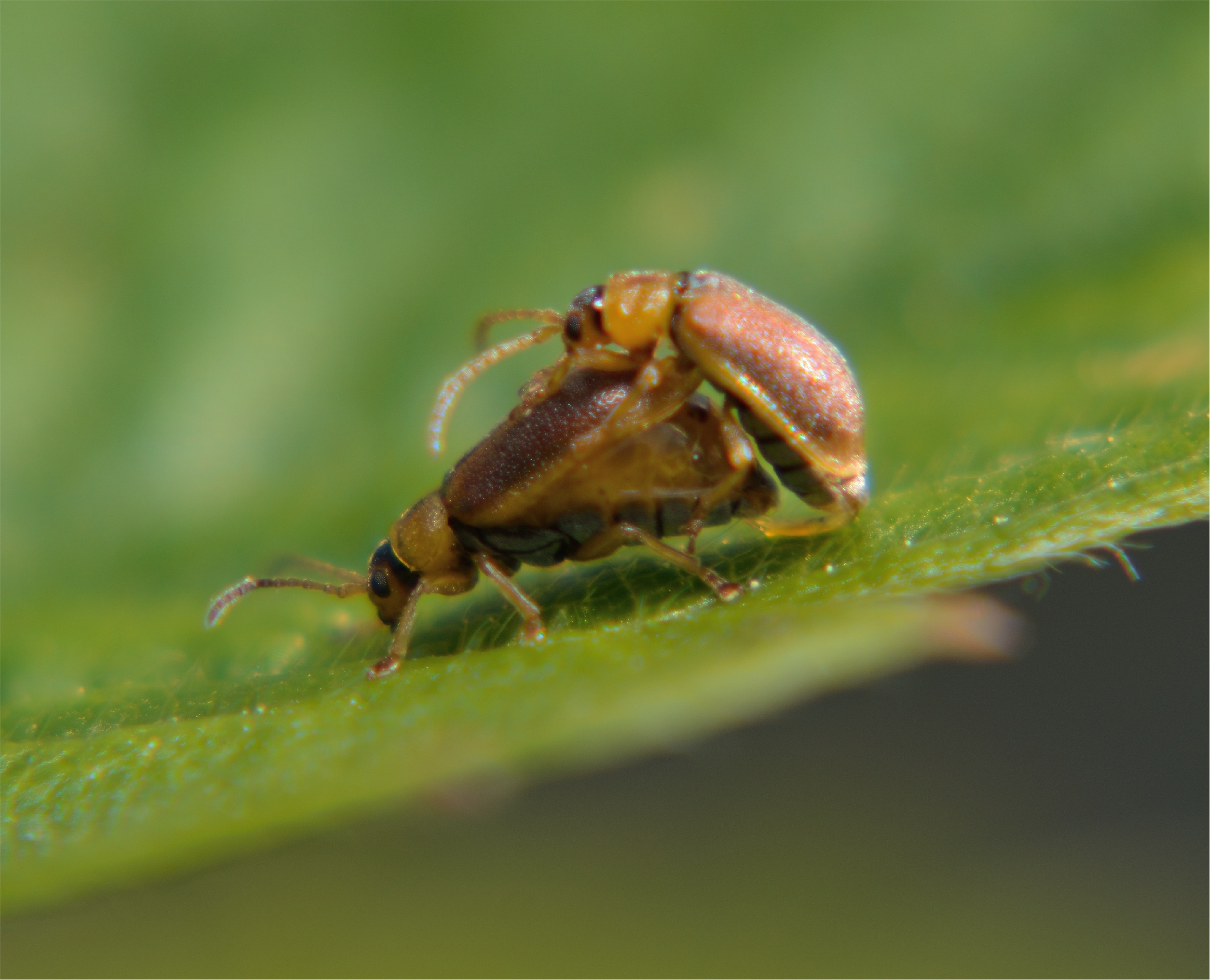
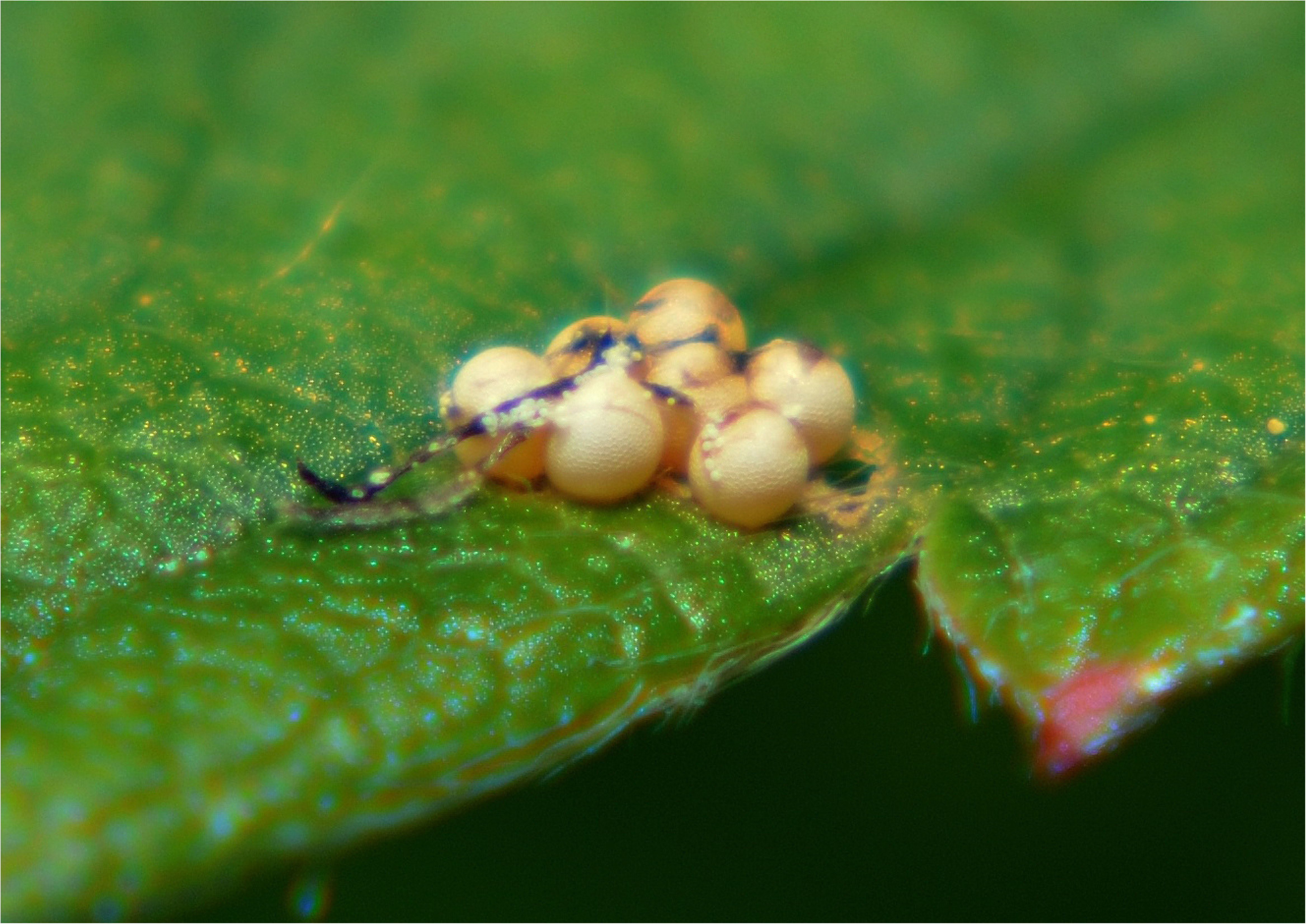
On the left, copulating strawberry leaf beetles, *Galerucella tenella*. On the right, eggs of *G. tenella* on strawberry leaf. Photo credits: Alejandro Ruete.

In order to facilitate a robust basis for sustainable pest management we here took both the antibiosis and antixenosis components of plant resistance into account (Stenberg and Muola 2017). Antibiosis has an adverse effect on insect herbivore performance, whereas antixenosis targets herbivore behaviour and renders the plant unattractive for feeding and/or oviposition (Painter 1951). To cover both aspects, we monitored oviposition preference as a proxy of antixenosis, as well as three proxies of antibiosis (egg hatching success, larval growth rate, and larval survival) for *G. tenella* in wild genotypes of *F. vesca*. In addition, we calculated the broad sense heritability of antibiosis (larval growth rate) in the screened *F. vesca* genotypes in order to investigate the phenotypic stability of this trait, with the potential view towards ‘rewilding’ strawberries. We hypothesized that: 1) wild *F. vesca* sampled within the native range of *G. tenella* in Sweden is likely to show high variation in the studied resistance parameters, and 2) that resistance should show significant broad-sense heritability, which would indicate its stability as a trait (i.e. that trait variation shows a significant genetic component) and hence breeding potential. Findings from this screening are likely to be important for the utilization of available genetic resources for restoring herbivore-resistance in cultivated strawberry (Andersen et al. 2015; Palmgren et al. 2015).

## Methods

### Study species

The woodland strawberry *Fragaria vesca* is a low-growing, herbaceous perennial plant. It occurs throughout most of the Holarctic including parts of Scandinavia and grows naturally in various half sunny and sunny habitats such as mountain slopes, roadsides, open forests, forest clearings and edges of forests and farmland (Roiloa & Retuerto 2007; Maliníková et al.2013; Schulze et al. 2012). The trifoliate leaves are evergreen, and it is an ever-bearing plant producing flowers and fruits throughout the entire growing season starting in May until late September (Maliníková et al.2013). Flowering peaks in early June and pollination of the white hermaphrodite flowers is carried out by insects, although selfing also commonly occurs alongside cross-pollination (Muola et al. 2017). In addition to the production of animal dispersed fruits, *F. vesca* is also capable of clonal reproduction and produces a high amount of runners that grow into self-sustaining plants in a short time (Schulze et al. 2012).

The oligophagous *Galerucella tenella* L. (Coleoptera: Chrysomelidae; Fig. 1) feeds on different species of the Rosaceae family, including *Fragaria* spp. (Stenberg et al. 2006; Hambäck et al. 2013). Wild meadowsweet (*Filipendula ulmaria* L.) serves as the main host, from which *G. tenella* commonly spills over to neighboring Rosaceae plants such as *Fragaria vesca*. *Galerucella tenella* has become a noticeable pest especially in organic strawberry cultivations. Outbreaks with severe damage have been reported from strawberry cultivations in the Nordic countries (Stenberg and Axelsson 2008; Stenberg 2012; Stenberg 2014), the Baltic States (Kaufmane and Libek 2000), and Russia (Bulukhto and Tsipirig 2004).

The adult *G. tenella* beetles hibernate in the topsoil and emerge during April to May. Mating as well as oviposition takes place directly on the host plant after emerging and the larvae hatch after few weeks from early June on depending on the temperatures in spring (Stenberg et al. 2006). Both adults and larvae feed on the leaves and flowers (Muola et al. 2017). Larvae might also tunnel into ripe berries and gnaw the fruit surface causing brown scaring [own observation, Parikka and Tuovinen 2014). After 2-4 weeks of feeding the larvae leave the host plant to pupate in the upper soil layer from early July on until mid-August in cold (Stenberg et al. 2006).

### Collection and experimental organization of wild germplasm

We adopted a quantitative genetic framework to study the genetic variation and heritability of resistance against *Galerucella tenella* using 86 wild *Fragaria vesca* genotypes. Following the protocol described by Muola et al. (2017) we collected wild plant genotypes from 86 randomly selected and geographically distinct locations (Fig. 2; see Online resource 1 for coordinates) across Uppsala County, Sweden in spring 2012. Uppsala County encompasses 8207 km^2^ and the distances between the sampling locations varied between 7 and 40 km. One genotype was collected from each site in order to maximize the geographic resolution of the sampled area. A recent microsatellite marker analysis where seven of these collected clones showed that they are true (separate) genotypes (Hilmarsson et al. 2017).

**Fig. 2.**
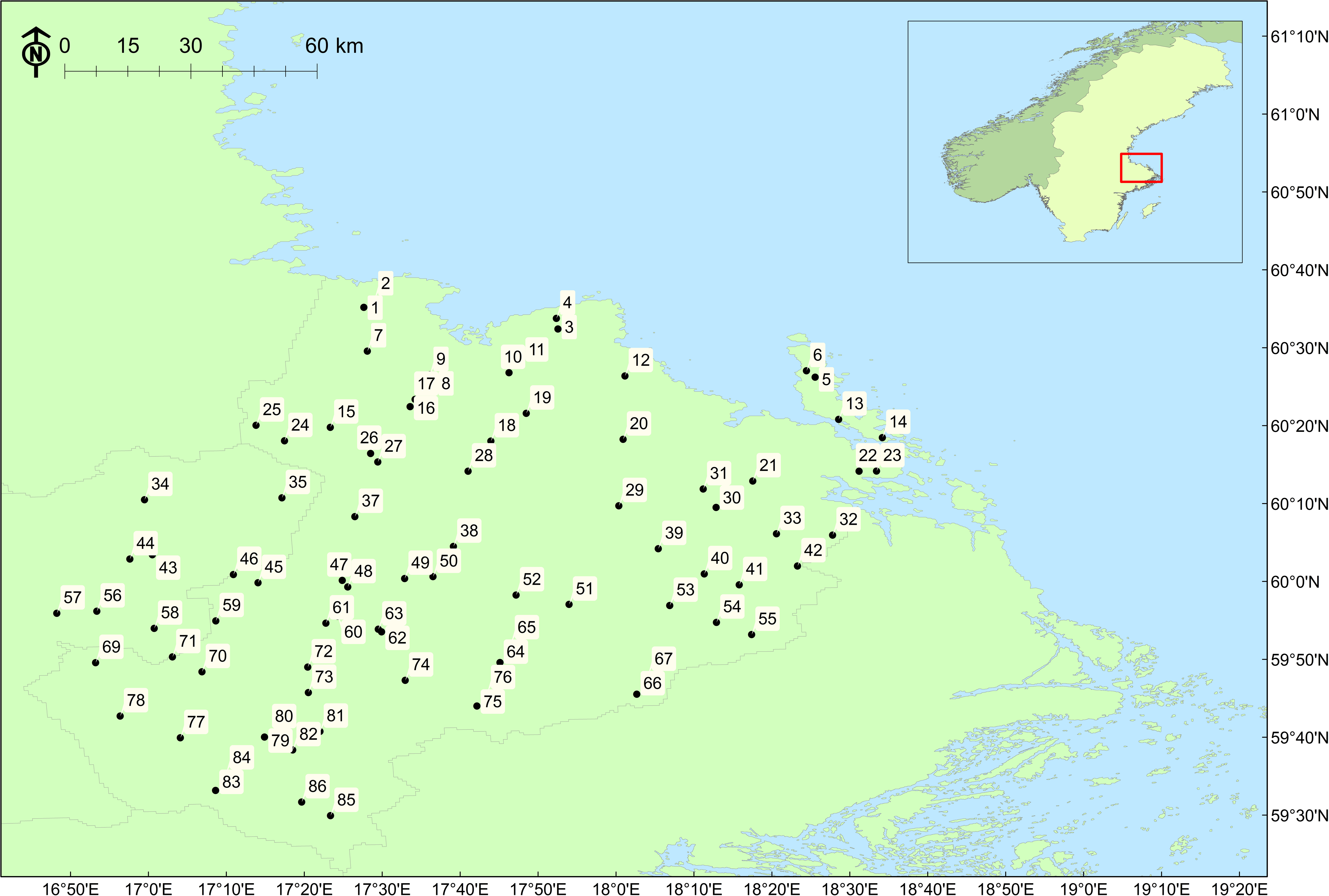
Map of the collection sites of 86 *Fragaria vesca* genotypes used in this study.

We cloned the sampled plant genotypes for several vegetative generations at the SLU Ultuna campus and planted 40 runners per plant genotype in blocks (one runner per plant genotype per block) in sandy soil in an open agricultural field (N59.741°, E17.684°), 15 km south of Uppsala in Autumn 2013. The distance between the plants was 50 cm and the entire common garden was covered with fabric mulch (Weibulls Horto) to reduce weed densities. The common garden was manually weeded when needed, but we did not apply any irrigation or fertilizer. The plants were growing in the common garden for two years before we used them in this study. The plant genotypes were specifically collected for the purpose of this study and corresponding plant material can be obtained from Stenberg on reasonable request.

### Screening of antibiosis

We measured antibiosis using several different proxies, namely larval survival, larval development time (days from hatching to pupation), pupal weigh (mg) and larval growth rate (mg/day, calculated from the two previous proxies). As growth rate takes both development time and weight into account we think it provides the most complete proxy of antibiosis. Low larval growth rate corresponds to high antibiosis and vice versa. As a final proxy for antibiosis we recorded the egg hatching success, which was done in line with the oviposition (antixenosis) screening and is therefore described under the antixenosis paragraph below.

For the screening of larval growth rate and survival we used detached leaves of the 86 plant genotypes that were collected from the common garden described above. To obtain neonate larvae for the experiment, we collected adult *Galerucella tenella* from four distinct natural meadowsweet populations in the vicinity of Uppsala (N59.809°, E17.667°; N59.788°, E17.664°; N59.806°, E17.652°; N59.806°, E17.665°) during early May 2015. The adult beetles were collected from locations with no co-occurring *Fragaria vesca* population to exclude a previous feeding experience with strawberry plants. This was done to avoid mistaking putative local adaptation of the herbivore with the genetic variation in plant resistance. The collected beetles were randomly mixed and placed in meshed cages containing a mix of *F. vesca* plants chosen randomly among the 86 genotypes. The cages were kept in a greenhouse (15°C, LD 16:8 h photoperiod, 80% RH) and the beetles were allowed to mate and oviposit freely on the offered plants. In order to rear the larvae, plants with eggs were retrieved and placed in a separate cage and monitored for hatching larvae.

We placed the neonate larvae (< 24h) individually in a 30 ml plastic containers and assigned them randomly to feed on one of the 86 plant genotypes. Ten larvae were assigned to each plant genotype, resulting in 860 rearing containers. We provided the larvae with one detached, undamaged, middle aged leaf from the assigned plant genotype. The leaf was replaced every third day and the rearing containers were cleaned in course with the leaf exchange. The rearing containers we kept in a climate chamber until the adult stage (15°C, LD 16:8 h photoperiod, 80% RH). We checked the larvae daily and weighed each larva on the day it reached the pupal stage. Furthermore, we recorded the survival and determined their sex after they reached the adult stage.

### Screening of antixenosis

We used oviposition preference as a proxy of antixenosis and addressed host plant suitability in the context of maternal choice. The oviposition preference of *Galerucella tenella* females was studied in a two-choice experiment. Based on the earlier larval growth rate screening (antibiosis, see description above) we selected eight plant genotypes with low larval growth rate (high antibiosis) and eight with high larval growth rate (low antibiosis) to test if ovipositing beetles discriminate between them. Runners from each of the 16 selected plant genotypes were collected from the common garden and propagated further to produce genetically identical replicates. The runners were potted in 0.2 liter pots with Hasselfors^TM^ (Hasselfors, Örebro, Sweden) planting soil and placed in a greenhouse (15°C, 16:8h) for five weeks during April 2016. The plants were then moved outside in early May 2016 one week prior to the experiment to allow them to adjust to the outside conditions. We used ten approximately equal-sized replicates from each plant genotype to exclude plant size as selection criteria for the female beetle, resulting in 80 replicates of both high and low antibiosis plants. The two types (high/low antibiosis) of plant genotypes were randomly paired and placed in meshed cages (40×40×40 cm) located at a sunny, sheltered place at the experimental garden of the SLU Alnarp campus. Within each cage, the two paired plants were spaced 30 cm apart. This was done to avoid leaf contact between the plants so that the female beetle had to choose one or the other. We collected the female *G. tenella* for the experiment from the same places as the beetles used in the performance experiment with two additional locations (N59.782°, E17.753°; N59.833°, E17.914°) during early May 2016 and selected mating couples with a gravid female to ensure subsequent egg laying. After collection the female beetles were placed centrally on the bottom of the cage and allowed to oviposit freely on the offered plant pair for 48h. For each plant, the number and the position (leaf blades or leaf petioles) of the laid eggs was noted. The position of the egg determines the microclimate for the egg development as well as the exposure of eggs and neonate larvae to predation or conspecific competition. Furthermore, the egg position facilitate or hamper the migration of the neonate larvae to young, developing leaves for feeding and hiding from predators.

After all eggs were counted, the plants were placed back into their cages to monitor the survival of the eggs as an additional aspect of antibiosis. Insect eggs can induce plant defences that harm the eggs (Hilker and Fatouros 2015). We measured the egg hatching success by allowing the eggs to hatch naturally and counting the first round of neonate larvae (<24 h) for each plant individual. Plants and *G. tenella* females were used only once in each oviposition choice trial.

### Statistical analyses

All the following data analyses were conducted with R, version 3.3.1 (R Core Team 2016). We used the inverse of herbivore performance measured as larval growth rate as one of three proxies for antibiosis in *Fragaria vesca* against *Galerucella tenella*. We analyzed the genetic variation in growth rate with a linear mixed model with larval growth rate as a response variable. Larval growth rate was calculated by dividing pupal weight (mg) by larval developmental time (days). The analyses of genetic variation in these two traits separately are presented in Online resource 2. We included plant genotype as a random factor and the sex of the beetles as a covariate to account for the potential sexual dimorphism in larval growth rate (or in pupal weight and development time). As per previous quantitative genetic analyses in *F. vesca* (Egan et al. 2018), this model structure – in which genotype is fitted as a random effect – permitted environmental (within-genotype) variation to be partitioned from genetic (between-genotype) variation (Hill 2010) in order to predict a ‘total genetic value’ for each genotype (Piepho et al. 2008). These values were used to visualize genetic variation in plots, or as an input to additional analyses, as detailed below. The interaction term between plant genotype and the covariate was insignificant and therefore removed from the final model. We assessed normality by visual examination and conducted a Levene’s test to check for equality of variances of the residuals.

As a second proxy of antibiosis we used one-way ANOVA to test whether larval growth rate (working with genotype genetic values as predicted in the previous model) is associated with larval survival rate, i.e. the percentage of larvae surviving per plant genotype. Due to the high rate of larval survival, we used the larval growth rate as response variable and grouped the survival rate into three categories of 100%, 90%, and less than 90% survival. This categorical factor was then used as an explanatory variable to avoid zero inflation in the model. We validated the model for normality and equality of variances as described above, and compared the survival levels in a posthoc comparison based on a Tukey’s pairwise analysis.

As a third proxy for antibiosis, we examined the effect of plant genotype on the survival of the eggs that were laid in the antixenosis screening (see below). As response variable we calculated the number of eggs successfully hatched as a proportion of the total amount of eggs per plant. We fitted the proportional response of egg hatching success in a generalized linear mixed model (logit link function) with a binomial error structure, and used antibiosis (i.e. larval growth rate) level (susceptible vs. resistant) as a fixed factor and individual plant genotype nested within the antibiosis level as a random factor. We performed a likelihood ratio test to assess the significance of the random effect of genotype. We validated the model with the Anderson Darling test and visual examination of the residuals.

To examine the antixenosis of *Fragaria vesca* genotypes we tested whether antibiosis affected the oviposition choice of *Galerucella tenella* females. Since the female beetles laid different absolute numbers of eggs on different plants, as well as on different tissues (i.e. the leaf blade and leaf petiole), we calculated the number of eggs laid on each tissue of a plant as a proportion of the total amount of eggs laid in the cage (following the set up in cages containing one resistant and one susceptible plant genotype). We used egg proportion per plant tissue (i.e. eggs laid on the leaf blade / all eggs laid in the cage, and, eggs laid on the leaf petiole / all eggs laid in the cage) as a proportional response variable in a binomial generalized mixed model (R-package “lme4”), in which plant antibiosis level (susceptible vs. resistant), plant tissue type, and their interaction were fitted as a fixed factor. Both plant individual nested within genotype, and genotype nested within the plant antibiosis level (susceptible vs. resistant) were included as random effects; in order to account for the non-independence of egg laying locations within the same plant individual, and the genotypic effect in the resistance level, respectively. Due to the significant interaction term between plant antibiosis level and plant tissue type, we obtained estimated marginal means and computed contrasts with the ‘emmeans’ package for R. As only four of the six possible combinations were of interest (e.g. resistant leaf blade vs. resistant leaf petiole; resistant leaf blade vs. susceptible leaf blade), we used the ‘holm’ method to adjust for four pairwise comparisons. To test whether the random effect of genotype had a significant effect, we ran a likelihood ratio test to compare the full model to a null model in which this random effect was removed. We validated the model through the Anderson Darling test for normality and visual examination of the residuals.

To examine broad-sense heritability (or clonal repeatability) in antibiosis (larval growth rate) among the tested plant genotypes, broad-sense heritability (*H*^2^) estimates were calculated from a linear mixed effects model. The tests for broad-sense heritability in larval development time and pupal weight separately are presented in Online resource 2. The model was fitted using the rptR package (Holger et al. 2016) in R (function ‘rpt’), and used the same model structure as specified for the resistance model above (i.e. ‘growth rate’ predicted by the random effect of ‘genotype’, controlling for the fixed effect of ‘sex’). The standard error of *H*^2^ was estimated based on 1000 parametric bootstraps, and the significance of *H*^2^ was tested via a likelihood ratio test.

### Results Antibiosis

We found genetic variation in antibiosis levels against *Galerucella tenella* in *Fragaria vesca* as indicated by statistically significant variation in the growth rate of *G. tenella* larvae reared on 86 different plant genotypes (χ^2^ =78.6, d.f.=1, P <0.001, Fig. 3). Furthermore, there was a heritable component to this variation, as broad-sense heritability in antibiosis (measured as larval growth rate) differed significantly from zero (*H*^2^ = 0.22 ± 0.04 SE). Similarly, both larval development time and pupal weight showed significant genetic variation between the tested plant genotypes and there was a heritable component in the observed variation (Online resource 2). The average larval growth rate varied from 0.204 ± 0.03 mg/day on the most susceptible plant genotype, to 0.129 ± 0.016 mg/day on the most resistant plant genotype. Female pupae were significantly heavier than male pupae (LMM: t-value =-13. 5, d.f.=733, P<0.001, Online resource 2) and, accordingly, females also had a significantly higher larval growth rate than males (F=139.52, d.f.=1, 717, P <0.001).

**Fig. 3.**
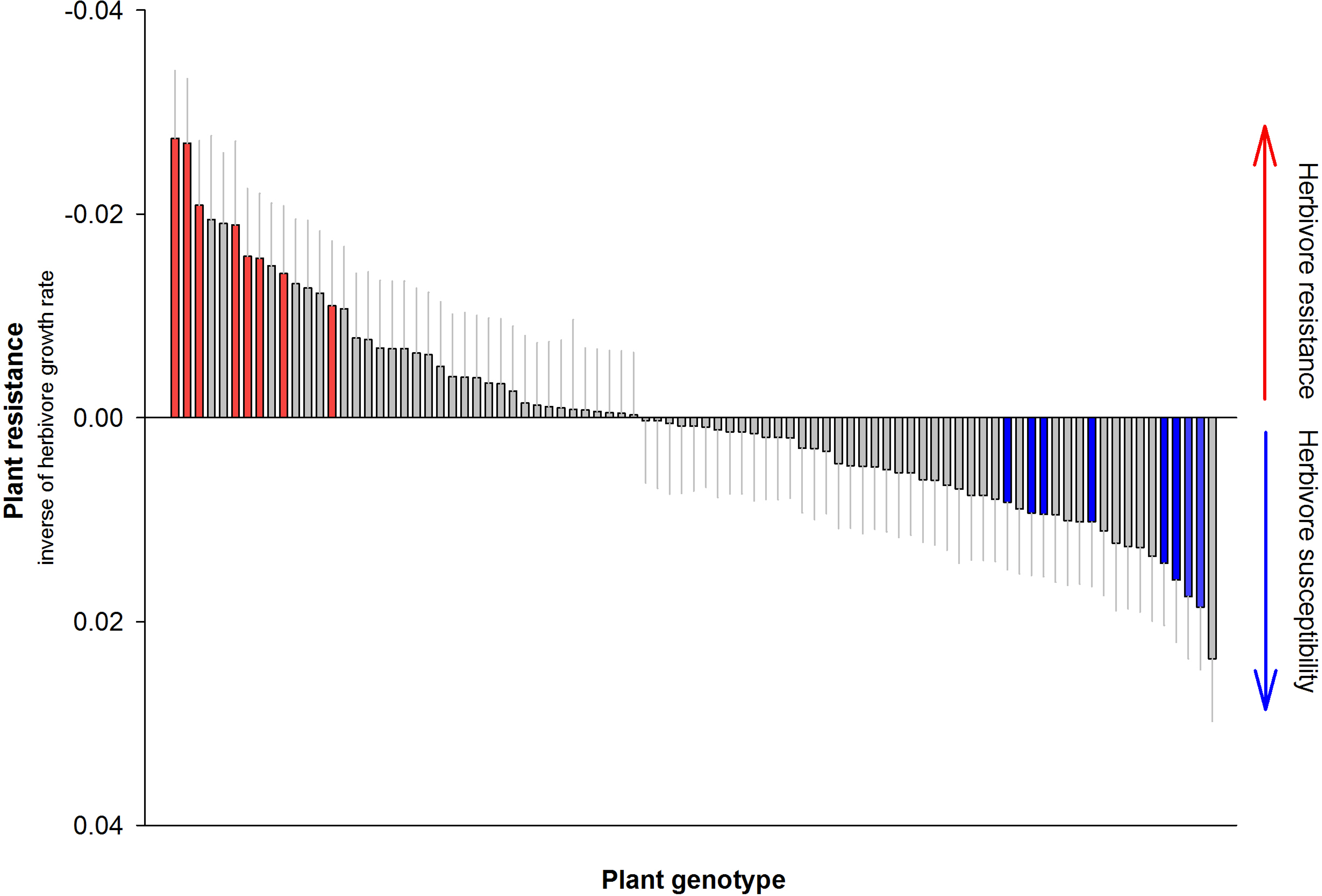
Genetic variation in plant resistance (antibiosis) against *Galerucella tenella* in 86 wild collected *Fragaria vesca* genotypes. Antibiosis was measured as inverse of herbivore performance. Larval growth rate (calculated as pupal weight divided by the larval development time from hatching until pupation) was used as a measure of herbivore performance. In the figure, the mean larval growth rate on each plant genotype is compared to the overall mean (standardized on zero) of all 86 plant genotypes used. Negative values i.e. larval growth rate lower than the overall mean indicate more resistant plant genotypes, while positive values indicate more susceptible plant genotypes. The red and blue bars denote the plant genotypes selected for experiments testing oviposition preference and egg survival. Predicted mean (total genetic value) + SE.

The overall survival rate of *Galerucella tenella* from egg hatching until pupation was high (93.2 %). However, larval growth rate and survival were associated: a higher than average larval growth rate was observed for those plant genotypes where all beetles survived, in contrast to below average growth rates observed in cases where one or more larvae died (F=9.63, d.f.=2, P<0.001; Fig. 4), indicating that antibiosis also affects survival through its effects on growth rate. Interestingly, egg hatching success was not affected by plant antibiosis level (Z=1.38; df= 160, P= 0.169; Fig. 5a), in which a predicted average of 47.4 % of eggs successfully hatched on a resistant genotype, and 56.5% on a susceptible genotype. However, hatching success was affected by plant genotype (χ^2^ =6.31, d.f.=1, P <0.012).

**Fig. 4.**
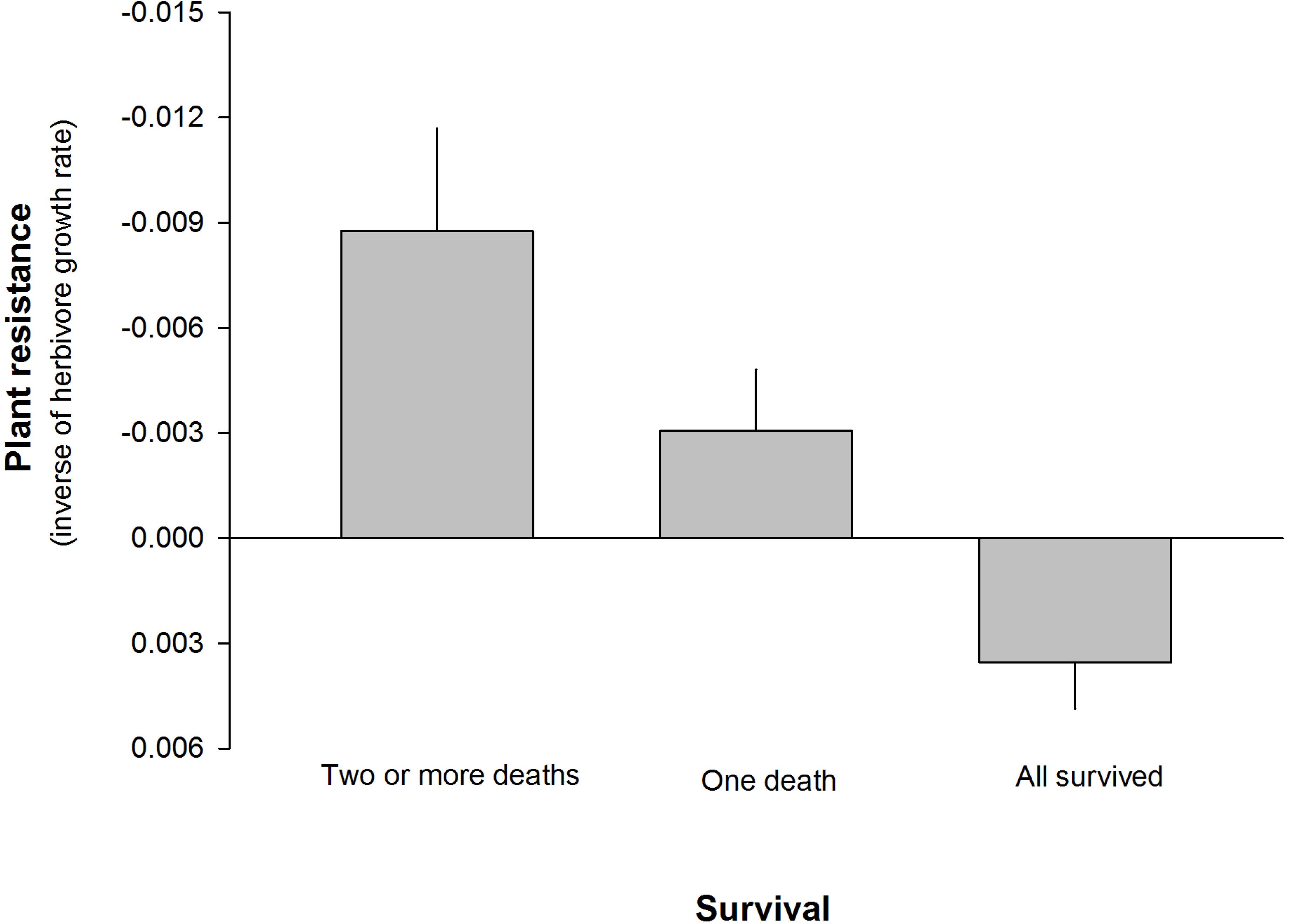
Association between larval growth rate and survival of *Galerucella tenella* larvae. The inverse of larval growth rate (calculated as pupal weight divided by the larval development time from hatching until pupation) was used as a proxy of plant resistance (antibiosis). Larval survival was divided into three classes according the number of larvae that survived when assigned to feed on the certain plant genotype. Statistical significance levels have been obtained by pairwise Tukey’s posthoc tests. Estimated marginal means + SE. ** 0.05 < *P* < 0.001; *** *P* <0.001.

**Fig. 5.**
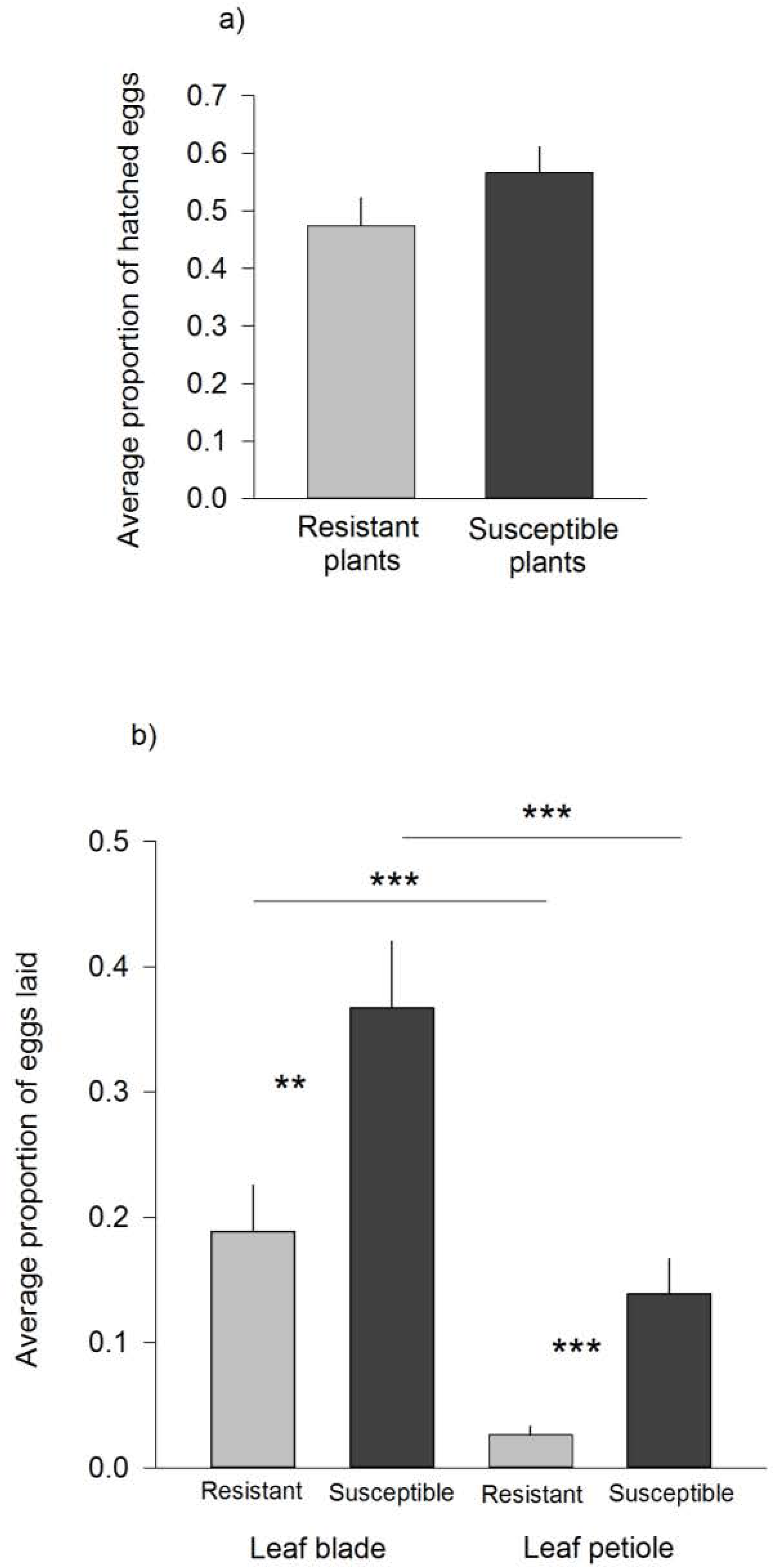
Effect of plant resistance (antibiosis) on a) egg hatching success (antibiosis), and b) the interactive effects of leaf tissue (leaf blade vs. leaf petiole) and plant antibiosis level on oviposition preference (antixenosis) of *Galerucella tenella*. Plant resistance (antibiosis) was measured as inverse of herbivore performance i.e. larval growth rate calculated as pupal weight divided by the larval development time from hatching until pupation. Pairwise *a priori* contrast comparisons were conducted to compare differences between resistant leaf blade vs. resistant leaf petiole, susceptible leaf blade vs. susceptible leaf petiole, resistant leaf blade vs. susceptible leaf blade and resistant leaf petiole vs. susceptible leaf petiole. Estimated marginal means + SE. ** 0.05 < *P* < 0.001; *** *P* <0.001.

### Antixenosis

We used oviposition preference as a measure of antixenosis. We found that the location of eggs – whether located on the leaf blade or leaf petiole – was affected by antibiosis, as shown by a significant interaction between plant antibiosis level (resistant or susceptible) and leaf tissue type (leaf blade or leaf petiole; Z= 2.15, df=160, P=0.032, Fig. 5b). Female beetles oviposit on average 1.9 times more eggs on leaf blades and 5.3 times more on leaf petioles of susceptible plants compared to resistant plants. Overall, female beetles were 3.2 times more likely to lay eggs on plant genotypes classified as susceptible (averaged across different tissue types), compared to resistant plants (Z= 2.774, df=320, P=0.006; Fig. 5b). In addition to plant antibiosis level, oviposition preference was affected by plant genotype (χ^2^ =1.77, d.f.=1, P=0.036).

## Discussion

Theory suggests that relatively high genetic variation can be found even in small geographic areas (Thompson 2013; Egan et al. 2018). Concordantly, we found significant genetic variation in the measured resistance traits against the Northern European pest insect *Galerucella tenella* in our screening of wild woodland strawberry, *Fragaria vesca*. In our study, we observed a large gradient in resistance, ranging from susceptible to highly resistant, suggesting that an untapped genetic resource is available in wild germplasm from Nordic populations of this plant species. However, the sampled area represents only a small fraction of this species’ Eurasian and North American distribution, suggesting that even higher variation could be harnessed if germplasm from the whole distribution was screened.

In order to obtain robust resistance in plants against pest insects, plant traits underlying strong antibiosis as well as antixenosis should be identified and optimized simultaneously (Stenberg and Muola 2017). We scored antibiosis using three different proxies: egg hatching success, larval growth rate and larval survival. The herbivore growth rate varied markedly and significantly between the plant genotypes. Furthermore, it correlated positively with the larval survival; i.e. plant genotypes that supported higher herbivore growth rate also contributed to higher herbivore survival. Although the mortality was generally low, on certain genotypes the herbivore growth rate was significantly reduced which can further act as a sub-lethal plant defence by prolonging the window of vulnerability of *G. tenella* to natural enemy attack (Clancy and Price 1987; Williams 1991). In addition, both larval development time and pupal weight, i.e. the intrinsic components of larval growth rate, also show individually significant genetic variation (Online resource 2). The plant traits and mechanisms underlying these antibiosis processes are yet to be identified, but our findings suggest that true resistance traits (e.g. chemical or physical defences), rather than low nutritional content, underlie antibiosis in woodland strawberry (Haukioja et al. 1991). Egg hatching success was unaffected by plant antibiosis level.

Our study of antixenosis – measured as oviposition preference – showed that this aspect of plant resistance also varied between plant genotypes. The female beetles showed a clear preference for genotypes with high host plant quality (low antibiosis), and thus by extension, an ability to make optimal choices for their offspring (*cf.* the mother knows best hypothesis) (Jaenike 1978; Valladares and Lawton 1991). Furthermore, this result connects the antixenosis component of *G. tenella* resistance in *F. vesca* positively with the antibiosis component. The fact that antibiosis and antixenosis correlate with each other is also beneficial from a future breeding perspective, as this opens up the possibility to simultaneously optimize both aspects of resistance in the same plant, providing robust protection against the target herbivore (Stenberg & Muola 2017).

Reverse breeding or rewilding is one promising way to provide modern crops with new or improved traits – such as resistance – which may have been lost or degraded during domestication. Given the continuous need to develop resistant strawberry cultivars, rewilding would allow cultivation of strawberry with reduced input of chemical pesticides. In the case of antibiosis against *G. tenella* we found different levels of broad-sense heritability (or clonal repeatability; the stability or ‘repeatability’ of this trait amongst clonal replicates of the same genotype) depending on which proxy that was used. Larval development time showed the highest level of heritability, while larval growth rate showed moderate levels, and pupal weight low heritability (Online resource 2). Which of the three proxies that is biologically most relevant is up to debate, and all of them have previously been used and recommended for other plant species (Stenberg and Muola 2017).

The genetic diversity of wild *Fragaria* species has previously provided novel genetic resources that were integrated into new cultivars (Liston et al. 2014). Among the wild *Fragaria* species, the woodland strawberry *F. vesca* is of special interest since its genome is sequenced (Shulaev et al. 2011) and displays a high level of collinearity with the cultivated garden strawberry in comparative mapping studies (Rousseau-Gueutin et al. 2008). New molecular methods such as CRISPR/Cas (Fonfara et al. 2016) and advances in plant breeding may soon offer solutions to overcome the multi gene nature of many resistance mechanisms and the differences in ploidy between wild and cultivated strawberry, and thus smooth the way for introgression of desired traits from wild relatives into modern crop varieties (Borges et al. 2018; Martinez et al. 2018).

## Conclusion

In conclusion, this study contributes to a growing body of literature showing that crop wild relatives in general, and woodland strawberry in particular, contain genetic resources that could contribute to resistance breeding in modern crops. In particular, we demonstrated considerable genotypic variation in each measured proxy of antibiosis (egg hatching success, larval growth rate and larval survival) and antixenosis (oviposition preference) as components of plant resistance against the pest insect *Galerucella tenella*. In terms of their combined potential for pest suppression, antibiosis and antixenosis either held no specific relation to each other, or else showed good compatibility – e.g. where genotypes which possessed high antixenosis levels against *G. tenella* larvae furthermore tended to be avoided by female adults as oviposition host plants. These results together highlight the large potential value of wild germplasm for resistance breeding. If breeding challenges can be overcome, we expect that “wild” resistance traits will play important role in future re-wilded strawberry cultivars, and contribute to a more sustainable crop production with reduced dependency on pesticides.

## Acknowledgements

We thank Joubin Haji Mirza Ali, Linn Holmstedt, Sara Janbrink, Wera Kleve, Joel Lönnqvist, Milda Norkute, Erika Qvarfordt, and Brenda Vidal Estévezfor technical assistance and Jan-Erik Englund and Adam Flöhr for valuable statistical discussions. The authors are funded by The Swedish Research Council Formas (grant nos. 217 - 2014-541 and 216 - 00223) and Maj and Tor Nessling Foundation (grant no. 201800048).

## Availability of data and material

Until publication, the data supporting our conclusions are available from the corresponding author on request. After publication data supporting our conclusions will be deposited to Dryad.

## Authors’ contributions

All authors conceived, organized and planned the research; JAS collected the wild accessions and established the common garden; DW undertook the experimental work and collected the data; DW, PAE, and AM analysed the data; DW led the writing of the manuscript. All authors contributed critically to the drafts and gave final approval for publication.

---

**Online resource 1 table 1.**
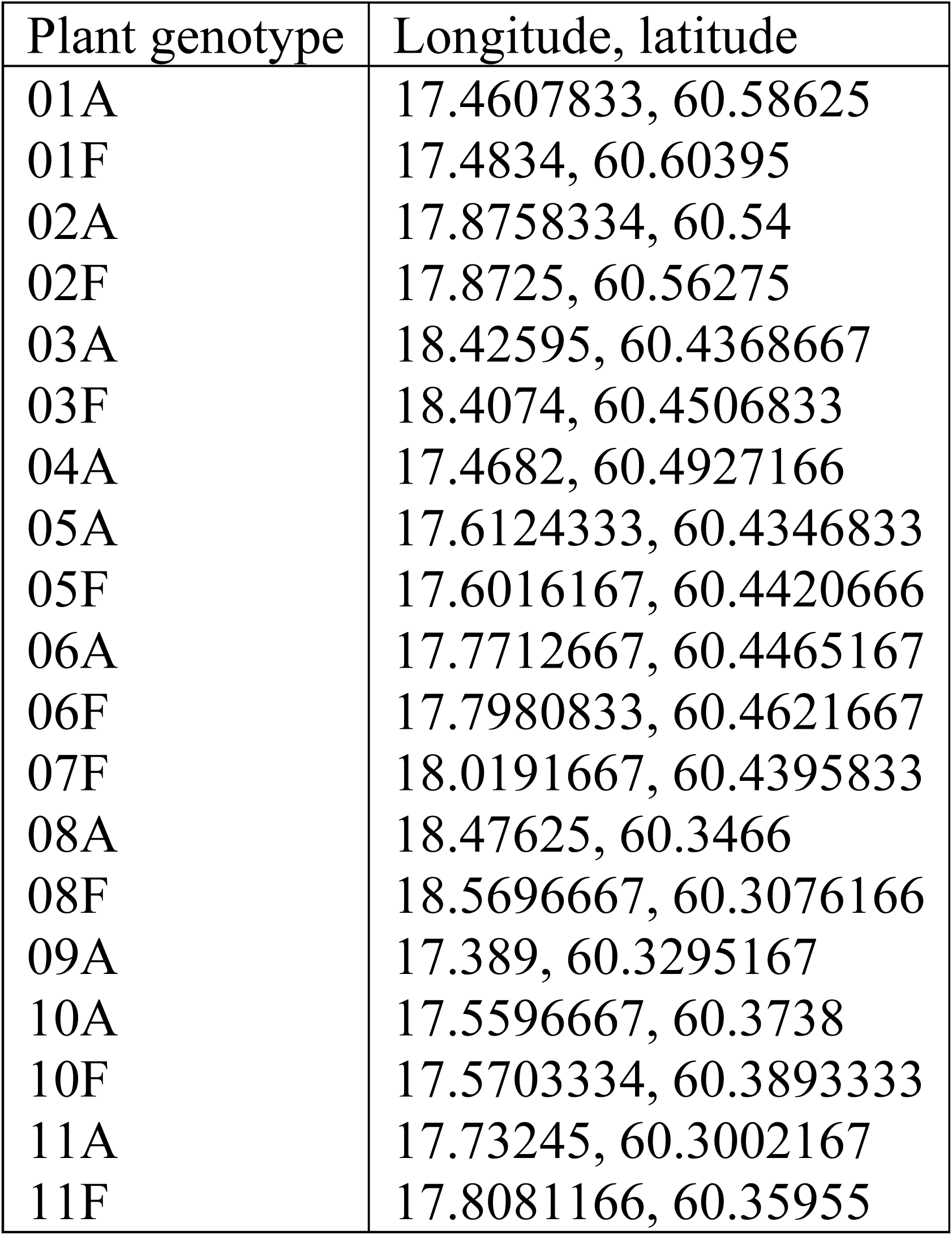
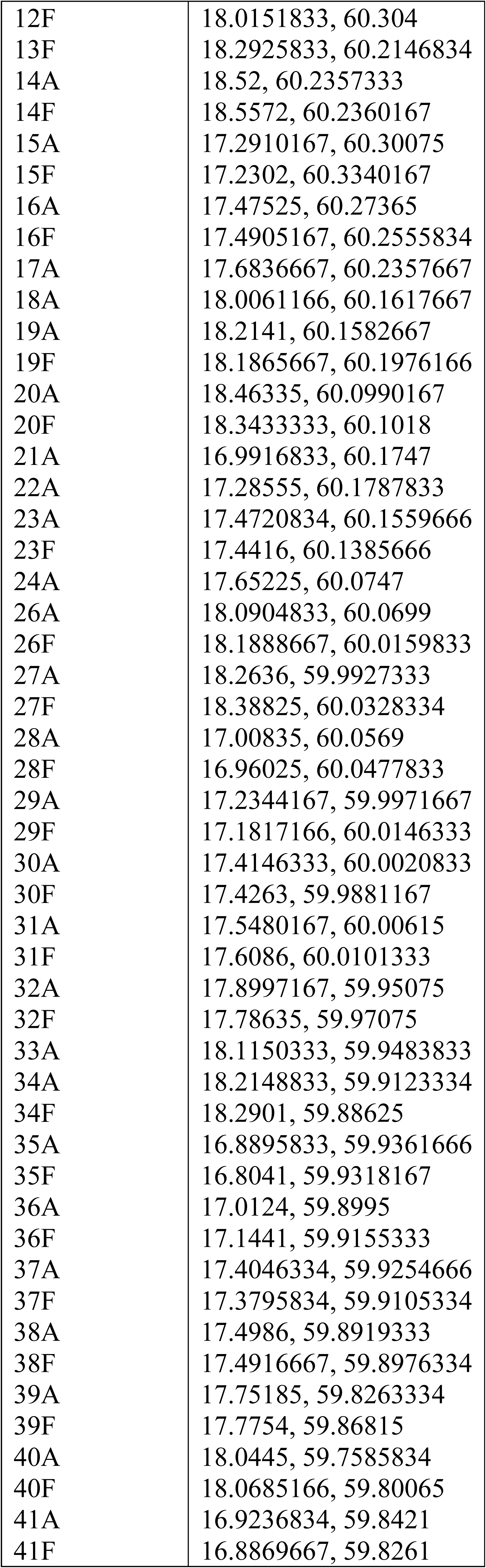
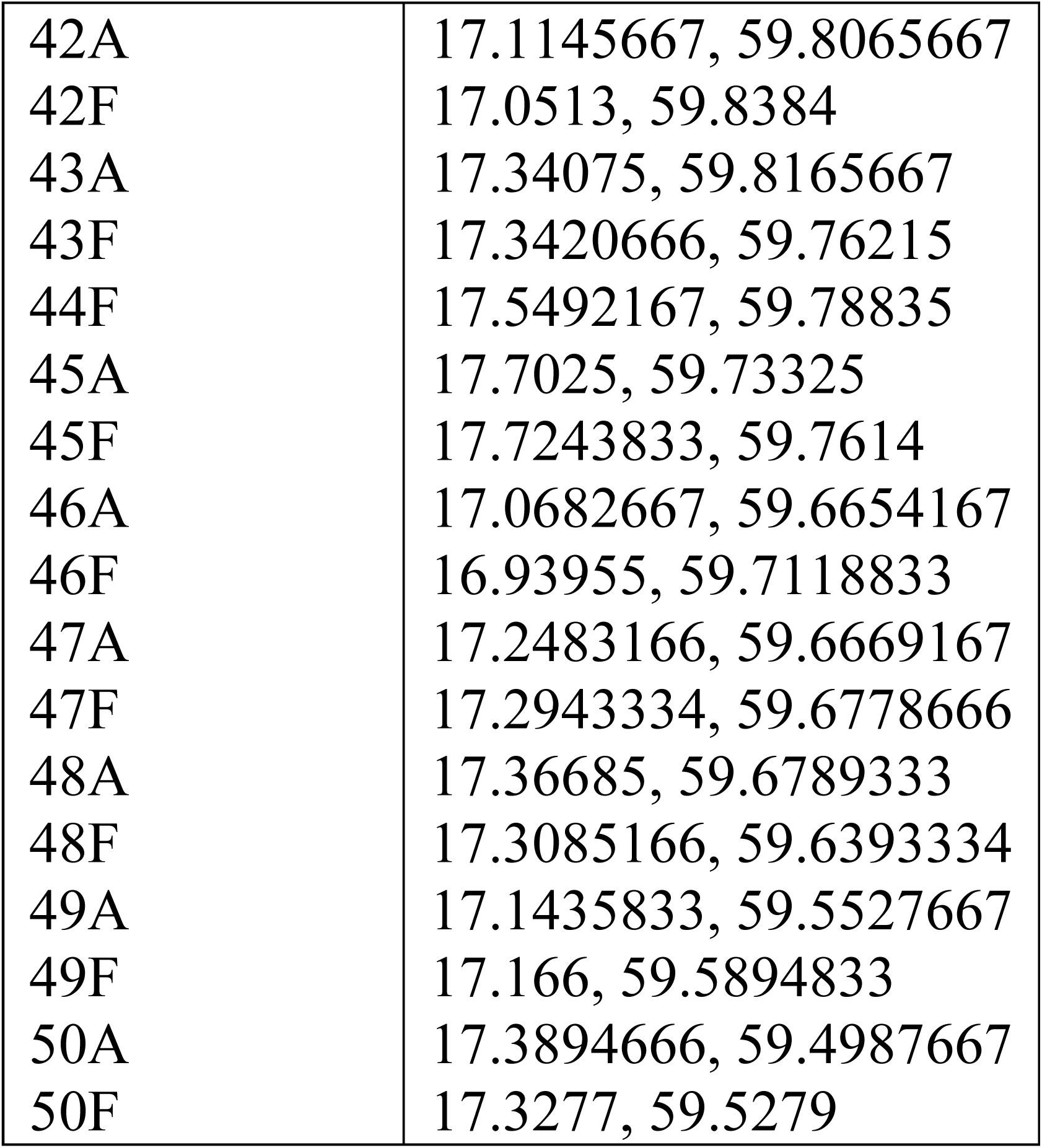
Collection site coordinates of woodland strawberry genotypes. Wild woodland strawberry genotypes used in the study were collected from 86 randomly selected and geographically distinct locations across Uppsala County, Sweden in spring 2012. Uppsala County encompasses 8207 km^2^ and the distances between the sampling locations varied between 7 and 40 km. Coordinates are given in decimal degree format and based on the international WGS 85 (World Geodetic System).

## Larval development time and pupal weight as proxies for antibiosis

We used 86 wild *Fragaria vesca* genotypes to study the genetic variation and heritability of plant resistance against *Galerucella tenella.* As one proxy for resistance, we used the inverse of herbivore performance measured as larval growth rate (i.e. antibiosis; reported in more details in the main text). Larval growth rate is composed of two measurements namely larval development time (days from hatching to pupation) and pupal weight (mg). Here, we report the results separately for these two traits.

We used the inverse of herbivore performance measured as larval development time and pupal weight as proxies for antibiosis in *F. vesca* against *G. tenella*. We analysed the genetic variation in larval development time and pupal weight with linear mixed model separately for each proxy. We included plant genotype as a random factor and the sex of the beetles as a covariate to account for the potential sexual dimorphism in larval development time or pupal weight. In addition, given the common inter-dependency between larval development time and pupal weight (i.e. that a larva needs to obtain a certain threshold weight before it can pupate), we included pupal weight as an additional covariate to the model analysing genetic variation in larval development time – and vice versa for the model analysing genetic variation in pupal weight. As such, these models hence quantified the extent to which unique (as opposed to overlapping) genetic variation existed for these traits. As per previous quantitative genetic analyses in *F. vesca* (Egan et al. 2018), this model structure – in which genotype is fitted as a random effect – permitted environmental (within-genotype) variation to be partitioned from genetic (between-genotype) variation (Hill 2010) in order to predict a ‘total genetic value’ for each genotype (Piepho et al. 2008). These values were used to visualize genetic variation in plots, or as an input to additional analyses, as detailed below. The interaction term between plant genotype and the covariate was insignificant and therefore removed from the final model. Larval development time did not fulfil the assumptions of normally distributed data, and, thus, we used the log-transformation. We assessed normality by visual examination and conducted a Levene’s test to check for equality of variances of the residuals.

To examine broad-sense heritability (or clonal repeatability) in larval development time and pupal weight among the tested plant genotypes, broad-sense heritability (*H*^2^) estimates werecalculated from a linear mixed effects model. The models were fitted using the rptR package (Holger et al. 2016) in R (function ‘rpt’), and used the same model structure as specified for the ‘larval development time’ and ‘pupal weight’ models above. The standard error of *H*^2^ was estimated based on 1000 parametric bootstraps, and the significance of *H*^2^ was tested via a likelihood ratio test.

We found genetic variation in antibiosis levels against *G. tenella* in *F. vesca* indicated by the statistically significant variation in both the development time (χ^2^ = 161.66, d.f.= 1, P<0.001) and pupal weight (χ^2^ = 25.01, d.f.= 1, P<0.001) of *G. tenella* larvae reared on 86 different plant genotypes (Supplementary Fig. 1a & b). The average larval development time varied from 16.8 days ± 1.0 SE on the most susceptible plant genotype, to 22.0 days ± 1.0 SE on the most resistant plant genotype. Pupal weight and larval development time were significantly related (LMM: t-value= -3.56, d.f.=741, P<0.001), in which larval development shortened by a predicted 0.02 days for every mg increase in pupal weight. Avarage pupal weight varied from 4.84 mg ± 0.11 SE when feeding on the leaves of the most susceptible plant genotypes to 3.82 mg ± 0.16 SE when feeding on the most resistant plant genotypes. Larval development time and pupal weight were significantly related (LMM: t-value= -4.31, d.f.=654, P<0.001), in which pupal weight was a predicted 0.04 mg lighter for every day’s increase in development time. Female pupae (4.46 mg ± 0.03 SE) were significantly heavier than male pupae (3.99 mg ± 0.03 SE) (LMM: t-value =-13. 5, d.f.=733, P<0.001). Furthermore, there was a heritable component in this variation, as broad-sense heritability both in larval development time and pupal weight differed significantly from zero (*H*^2^ = 0.36 ± 0.046 SE for larval development time, and *H*^2^ =0.112 ± 0.031 SE for pupal weight).

**Online Resource 2 Figure 1.**
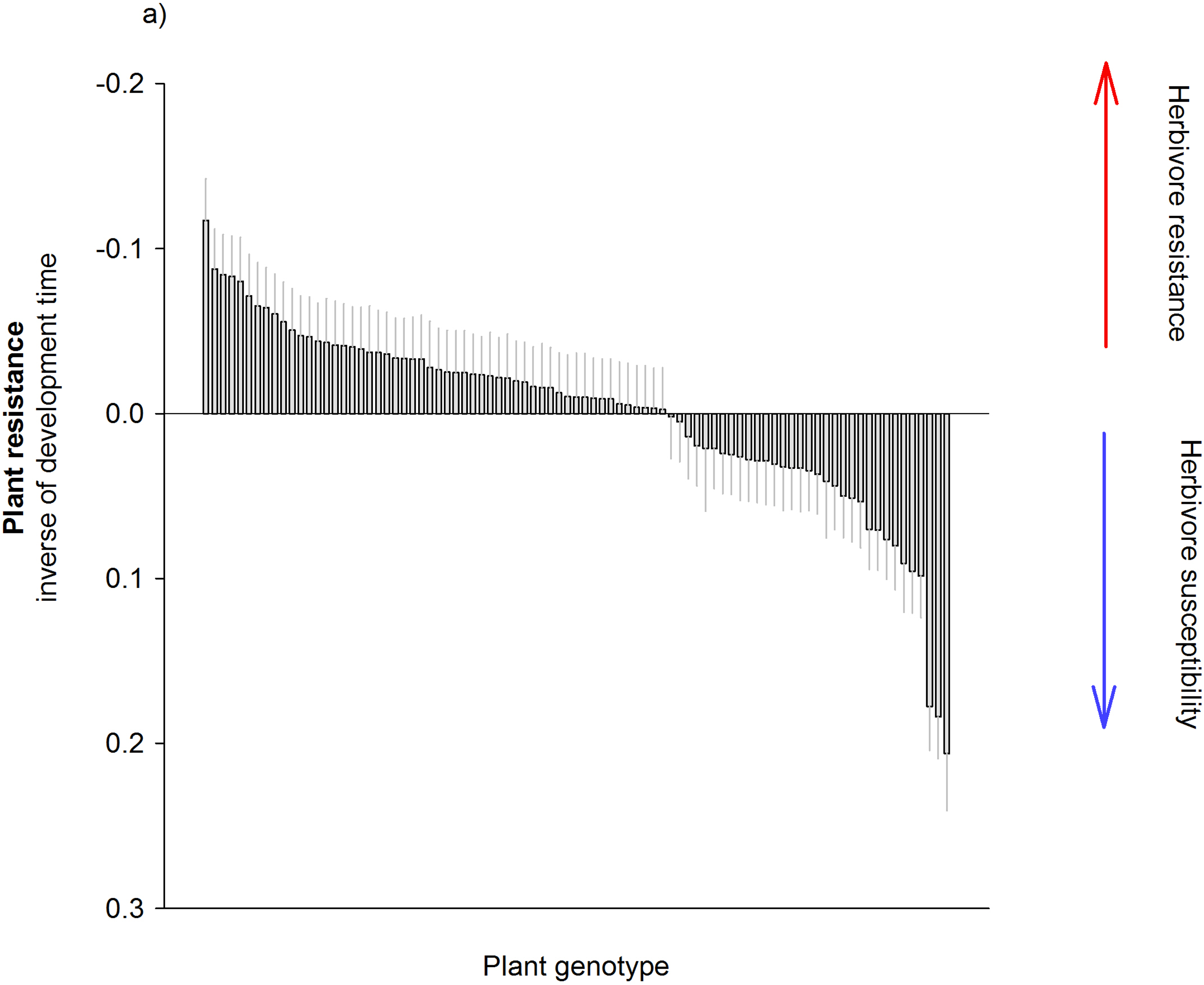
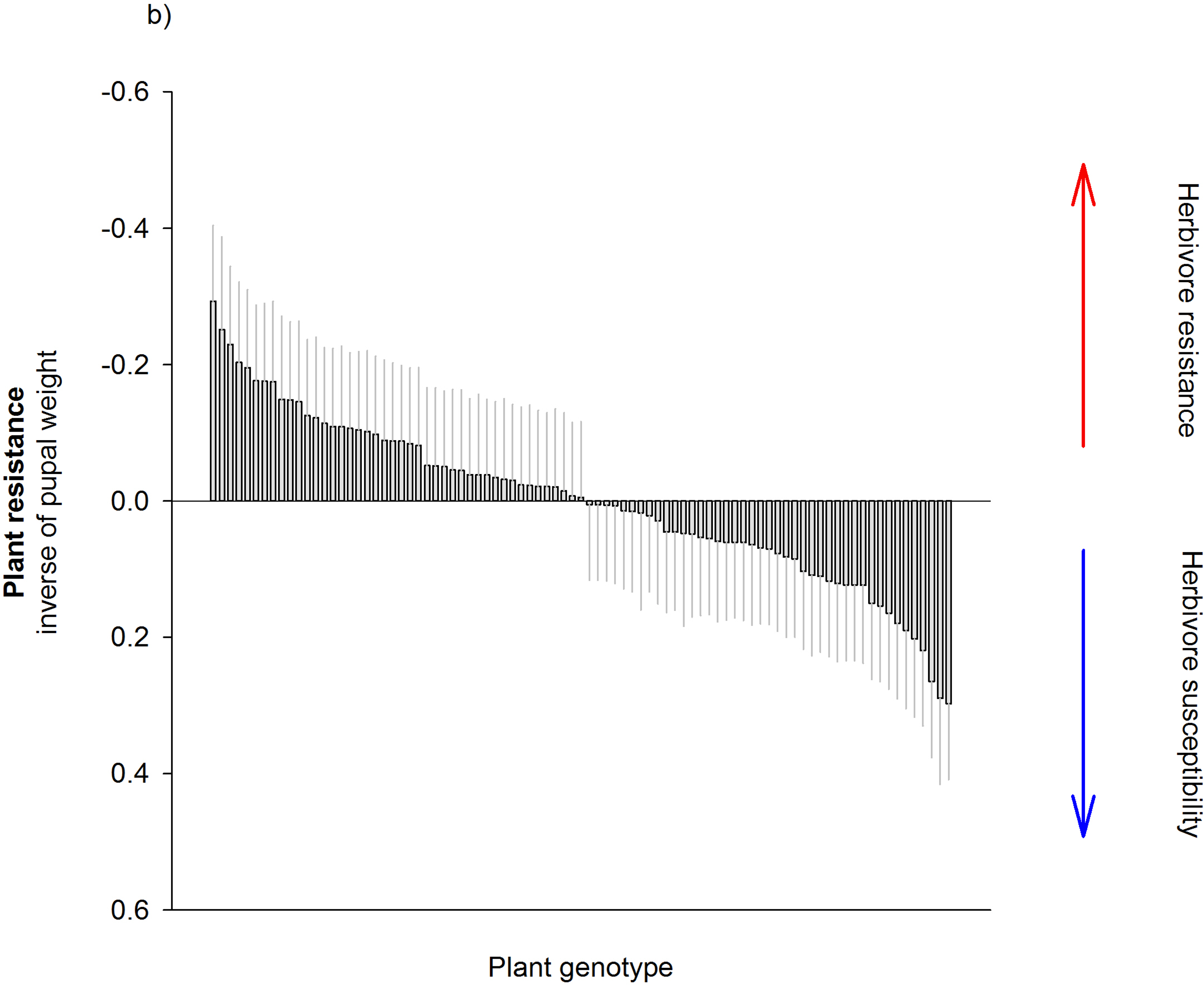
Genetic variation in plant resistance (antibiosis) against *Galerucella tenella* in 86 wild collected *Fragaria vesca* genotypes. Antibiosis was measured as inverse of herbivore performance. Larval development time from hatching until pupation (1a) and pupal weight (mg; 1b) were used as measures of herbivore performance. In the figure, the mean development time (1a) and the mean pupal weight (1b) on each plant genotype is compared to the overall mean (standardized on zero). Negative values indicate more resistant plant genotypes, while positive values indicate more susceptible plant genotypes. Predicted mean (total genetic value) + SE.

## References

Amil-Ruiz F, Blanco-Portales R, Muñoz-Blanco J, Caballero JL (2011) The strawberry plant defense mechanism: A molecular review. Plant Cell Physiol 52:1873–1903.

Andersen MM, Landes X, Xiang W, Anyshchenko A, Falhof J, Østerberg JT, Olsen LI, Edenbrandt AK, Vedel SE, Thorsen BJ, Sandøe P, Gamborg C, Kappel K, Palmgren MG (2015) Feasibility of new breeding techniques for organic farming. Trends Plant Sci 20:426–434.

Borges F, Parent J-S, van Ex F, Wolff P, Martínez G, Köhler C, Martienssen RA (2018) Transposon-derived small RNAs triggered by miR845 mediate genome dosage response in *Arabidopsis*. Nat Genet 50:186–192.

Broekgaarden C, Snoeren TAL, Dicke M, Vosman B (2011) Exploiting natural variation to identify insect-resistance genes. Plant Biotechnol J 9:819–825.

Bulukhto NP, Tsipirig OV (2004) Strawberry leaf beetle. Zashchita i Karantin Rastenii 8:42–43.

Chen YH, Gols R, Benrey B (2015) Crop domestication and naturally selected species interactions. Annu Rev Entomol 60:35–58.

Clancy KM, Price PW (1987) Rapid herbivore growth enhances enemy attack - sublethal plant defenses remain a paradox. Ecology 68:733–737.

Dempewolf H, Baute G, Anderson J, Kilian B, Smith C, Guarino L (2017) Past and future use of wild relatives in crop breeding. Crop Sci 57:1070–1082.

Doumett S, Fibbi D, Cincinelli A, Giordani E, Nin S, Del Bubba M (2011) Comparison of nutritional and nutraceutical properties in cultivated fruits of *Fragaria vesca* L. produced in Italy. Food Res Int 44:1209–1216.

Egan PA, Muola A, Stenberg JA (2018) Capturing genetic variation in crop wild relatives: An evolutionary approach. Evol Appl 11:1293–1304.

Ekuere UU, Dosdall LM, Hills M, Keddie AB, Kott L, Good A (2005) Identification, mapping, and economic evaluation of QTLs encoding root maggot resistance in *Brassica*. Crop Sci 45:371–378.

Fernandes VC, Domingues VF, Mateus N, Delerue-Matos C (2011) Organochlorine pesticide residues in strawberries from integrated pest management and organic farming. J Agric Food Chem 59:7582–7591.

Fonfara I, Richter H, Bratovič M, Le Rhun A, Charpentier E (2016) The CRISPR-associated DNA-cleaving enzyme Cpf1 also processes precursor CRISPR RNA. Nature 532:517–521.

Godfray HCJ, Blacquière T, Field LM, Hails RS, Petrokofsky G, Potts SG, Raine NE, Vanbergen AJ, McLean AR (2014) A restatement of the natural science evidence base concerning neonicotinoid insecticides and insect pollinators. Proc R Soc B 281:20140558.

Hambäck PA, Weingartner E, Ericson L, Fors L, Cassel-Lundhagen A, Stenberg JA, Bergsten J (2013) Bayesian species delimitation reveals generalist and specialist parasitic wasps on *Galerucella* beetles (Chrysomelidae): sorting by herbivore or plant host? BMC Evol Biol 13:92.

Haukioja E, Ruohomäki K, Suomela J, Vuorisalo T (1991) Nutritional quality as a defense against herbivores. For Ecol Manag 39:237–245.

Hilker M, Fatouros NE (2015) Plant responses to insect egg deposition. Annu Rev Entomol 60:493–515.

Hill WG (2010) Understanding and using quantitative genetic variation. Philos Trans R Soc Lond B Biol Sci 365:73–85.

Hilmarsson HS, Hytönen T, Isobe S, Göransson M, Toivainen T, Hallsson JH (2017) Population genetic analysis of a global collection of *Fragaria vesca* using microsatellite markers. PLoS ONE 12: e0183384.

Holger S, Stoffel M, Nakagawa S (2016) rptR: Repeatability Estimation for Gaussian and Non-Gaussian Data. R package version 0.9.0. 2016. https://CRAN.R-project.org/package=rptR Accessed 14 Aug 2018.

Hollender CA, Kang C, Darwish O, Geretz A, Matthews BF, Slovin J, Alkharouf N, Liu Z (2014) Floral transcriptomes in woodland strawberry uncover developing receptacle and anther gene networks. Plant Physiol 165:1062–1075.

Jaenike J (1978) On optimal oviposition behavior in phytophagous insects. Theor Popul Biol 14:350–356.

Kang C, Darwish O, Geretz A, Shahan R, Alkharouf N, Liu Z (2013) Genome-scale transcriptomic insights into early-stage fruit development in woodland strawberry *Fragaria vesca*. Plant Cell 25:1960–1978.

Kaufmane E, Libek A (2000) The occurrence of arthropods in strawberry plantation depending on the method of cultivation. Proceedings of the International Conference Fruit Production and Fruit Breeding Tartu 204–208.

Liston A, Cronn R, Ashman T-L (2014) *Fragaria*: A genus with deep historical roots and ripe for evolutionary and ecological insights. Am J Bot 101:1686–1699.

Longhi S, Giongo L, Buti M, Surbanovski N, Viola R, Velasco R, Ward JA, Sargent DJ (2014) Molecular genetics and genomics of the Rosoideae: State of the art and future perspectives. Hortic Res 1:1.

Malchev I, Fletcher R, Kott L (2010) Breeding of rutabaga (*Brassica napus* var. *napobrassica* L. Reichenb.) based on biomarker selection for root maggot resistance (*Delia radicum* L.). Euphytica 175:191–205.

Maliníková E, Kukla J, Kuklová M, Balážová M (2013) Altitudinal variation of plant traits : morphological characteristics in *Fragaria vesca* L. (Rosaceae). Ann For Res 56:79–89.

Martinez G, Wolff P, Wang Z, Moreno-Romero J, Santos-González J, Conze LL, DeFraia C, Slotkin RK, Köhler C (2018) Paternal easiRNAs regulate parental genome dosage in *Arabidopsis*. Nat Genet 50:193–198.

Mouhu K, Kurokura T, Koskela EA, Albert VA, Elomaa P, Hytönen T (2013) The *Fragaria vesca* homolog of SUPPRESSOR OF OVEREXPRESSION OF CONSTANS1 represses flowering and promotes vegetative growth. Plant Cell 25:3296–3310.

Muola A, Weber D, Malm LE, Egan PA, Glinwood R, Parachnowitsch AL, Stenberg JA (2017) Direct and pollinator-mediated effects of herbivory on strawberry and the potential for improved resistance. Front Plant Sci 8:823.

Olofsson E, Pettersson ML (1992) Leaf beetles in the genus *Galerucella* (Col., Chrysomelidae) on strawberry. Växtskyddsnotiser 56:42–45.

Painter RH (1951) Insect Resistance in Crop Plants. The University Press of Kansas, Lawrence.

Palmgren MG, Edenbrandt AK, Vedel SE, Andersen MM, Landes X, Østerberg JT, Falhof J, Olsen LI, Christensen SB, Sandøe P, Gamborg C, Kappel K, Thorsen BJ, Pagh P (2015) Are we ready for back-to-nature crop breeding? Trends Plant Sci 20;155–164.

Parikka P, Tuovinen T (2014) Plant protection challenges in strawberry production in northern Europe. Acta Hortic 1049:173–179.

Piepho HP, Moehring J, Melchinger AE, Buechse A (2008) BLUP for phenotypic selection in plant breeding and variety testing. Euphytica 161:209–228.

R Core Team (2016) R: A language and environment for statistical computing. R Foundation for Statistical Computing. Vienna.

Roiloa SR, Retuerto R (2007) Responses of the clonal *Fragaria vesca* to microtopographic heterogeneity under different water and light conditions. Environ Exp Bot 61:1–9.

Rousseau-Gueutin M, Lerceteau-Köhler E, Barrot L, Sargent DJ, Monfort A, Simpson D, et al (2008) Comparative genetic mapping between octoploid and diploid *Fragaria* species reveals a high level of colinearity between their genomes and the essentially disomic behavior of the cultivated octoploid strawberry. Genetics 179:2045–2060.

Shulaev V, Sargent DJ, Crowhurst RN, Mockler TC, Folkerts O, Delcher AL, et al (2011) The genome of woodland strawberry (*Fragaria vesca*). Nature Genet 43:109–116.

Schulze J, Rufener R, Erhardt A, Stoll P (2012) The relative importance of sexual and clonal reproduction for population growth in the perennial herb *Fragaria vesca*. Popul Ecol 54:369– 380.

Smith CM, Clement SL (2012) Molecular bases of plant resistance to arthropods. Annu Rev Entomol 57:309–328.

Stenberg JA (2012) Effects of local vegetation and plantation age for the parasitoid *Asecodes mento* – a biocontrol agent in organic strawberry fields. Insect Sci 19:604–608.

Stenberg JA (2014) Density-dependent herbivory and biocontrol of the strawberry leaf beetle, *Galerucella tenella*. Acta Hortic 1049:647–650.

Stenberg JA, Axelsson EP (2008). Host race formation in the meadowsweet and strawberry feeding leaf beetle *Galerucella tenella*. Basic Appl Ecol 9:260–267.

Stenberg JA, Muola A (2017) How should plant resistance to herbivores be measured? Front Plant Sci 8:663.

Stenberg JA, Witzell J, Ericson L (2006) Tall herb herbivory resistance reflects historic exposure to leaf beetles in a boreal archipelago age-gradient. Oecologia 148:414–425.

Tennessen JA, Govindarajulu R, Ashman T-L, Liston A (2014) Evolutionary origins and dynamics of octoploid strawberry subgenomes revealed by dense targeted capture linkage maps. Genome Biol Evol 6:3295–3313.

Thompson JN (2013) Relentless Evolution. University of Chicago Press, Chicago.

Ulrich D, Komes D, Olbricht K, Hoberg E (2007) Diversity of aroma patterns in wild and cultivated *Fragaria* accessions. Genet Resour Crop Evol 54:1185–1196.

Urrutia M, Schwab W, Hoffmann T, Monfort A (2015) Genetic dissection of the (poly)phenol profile of diploid strawberry (*Fragaria vesca*) fruits using a NIL collection. Plant Sci 242:151– 168.

Valladares G, Lawton JH (1991). Host-plant selection in the holly leaf-miner: does mother know best? J Anim Ecol 60:227–240.

## References

Holger S, Stoffel M, Nakagawa S (2016) rptR: Repeatability Estimation for Gaussian and Non-Gaussian Data. R package version 0.9.0. 2016. https://CRAN.Rproject.org/package=rptR Accessed 14 Aug 2018.

Williams IS (1991) Slow-growth, high-mortality – a general hypothesis, or is it? Ecol Entomol 24:490–495.

